# Profile of the somatic mutational landscape in breast tumors from Hispanic/Latina women

**DOI:** 10.1101/2022.08.01.502349

**Authors:** Yuan C Ding, Hanbing Song, Aaron Adamson, Daniel Schmolze, Donglei Hu, Scott Huntsman, Linda Steele, Carmina Patrick, Natalie Hernandez, Charleen D Adams, Laura Fejerman, Kevin L. Gardner, Anna María Nápoles, Eliseo J. Pérez-Stable, Jeffrey N. Weitzel, Henrik. Bengtsson, Franklin W. Huang, Susan L. Neuhausen, Elad Ziv

## Abstract

Breast cancer causes the most cancer deaths among Hispanic/Latinas (H/L). However, limited tumor-sequencing data from H/L are available to guide treatment. To address this gap, we performed whole-exome sequencing of DNA from 140 HL germline and 146 matched breast tumors and RNA-seq for the tumors. We generated somatic-mutation profiles, identified copy-number alterations (CNAs), and compared results to non-Hispanic White (White) women in The Cancer Genome Atlas. Similar to Whites, *PIK3CA* and *TP53* were the most commonly mutated genes in breast tumors from H/L. We found 4 common COSMIC mutation signatures (1, 2, 3, 13) and signature 16 not previously reported in other breast-cancer datasets. We observed recurrent amplifications in breast-cancer drivers including *MYC, FGFR1, CCND1*, and *ERBB2*, and a recurrent amplification on 17q11.2 associated with high *KIAA0100* gene expression, implicated in breast-cancer aggressiveness. Expanded research is required to determine how these characteristics of H/L tumors impact treatment response and survival.

## INTRODUCTION

Sequencing studies of breast cancer have identified recurrently mutated genes and somatic copy number alterations (SCNAs) affecting tumor suppressors and oncogenes^1-3^. Both somatic mutations and CNAs may be useful in determining prognosis. Currently, therapies for breast cancer can be selected based on particular somatic mutations (i.e., alpelisib for *PIK3CA*^4^), SCNAs (i.e., Trastuzumab for *HER2*), and germline mutations in genes in the homologous recombination repair (HRR) pathway (polyADPribose polymerase inhibitors -PARPi’s).

Genetic ancestry is associated with specific somatic mutations in many cancer types. *EGFR* mutations are approximately four-fold more common in lung cancer from women and men of East-Asian ancestry compared with lung cancer from women and men of other populations^5^ with self-reported Hispanic/Latinos (H/L) representing an intermediate group ^6, 7^. *FOXA1* mutations in prostate cancer also are substantially more common in East-Asian ancestry populations compared to European and African ancestry populations^8^. Comprehensive analyses of The Cancer Genome Atlas (TCGA) have demonstrated that many mutations and CNAs are more common in specific ancestral populations^9, 10^. In breast cancer, previous studies have demonstrated that women of African ancestry have higher rates of *TP53* mutations and lower rates of *PIK3CA* mutations, likely related to a higher incidence of a basal-like breast-cancer subtype in African-American women^11, 12^. However, the genomic landscape of breast cancer has not been well-characterized in H/L groups.

H/L represent the largest minority population in the US and have diverse origins, with the largest subpopulations including Mexican Americans and Puerto Ricans. Genetically, H/L are a population of mixed European, Indigenous-American (IA) and African ancestries with those ancestry proportions varying widely depending on country of origin and regions within a country. Although breast cancer is less common overall among H/L compared to self-reported non-Hispanic White (White) women due to both environmental ^13^ and genetic factors^14^, there is a higher proportion of breast cancers diagnosed under age 50 years than in Whites^15^. Moreover, outcomes are usually worse among H/L compared to White women^16^. In some studies, IA ancestry was associated with poorer outcomes among H/L with breast cancer^17^. Human epidermal growth factor (HER2) amplifications are over-represented among H/L and are more common among H/L with more IA ancestry compared to those with more European ancestry^18^. Few studies have investigated the distribution of somatic mutations and SCNAs in breast tumors from H/L. In TCGA, out of 1,096 breast-cancer cases, only 39 are self-reported H/L. A recent study analyzed data including whole-exome sequencing (WES) and gene expression data from 109 Mexican women living in Mexico ^19^. However, no similar-size study has been conducted in H/L in the United States (US). To investigate the somatic-mutational spectrum in breast cancer among H/L, we generated whole-exome sequencing (WES) and RNA sequencing (RNA-seq) data from 146 tumors from 140 H/L from Southern California and performed analyses of somatic mutations, SCNAs, and gene expression.

## METHODS

### Participants

One hundred and forty breast-cancer patients seen at City of Hope (COH) in Duarte, California were included in this study. All participants signed a written informed consent approved by the COH Institutional Review Board. Inclusion criteria were: 1) self-identified as H/L; 2) tumor tissue from surgery was available and the sample contained more than 40% tumor based on examination by a single breast pathologist (D. Schmolze). The percentage tumor ranged from 40% to 90% with an average of 64% and a median of 65% tumor. An exclusion criterion was neo-adjuvant therapy as treatment could change the mutation profile. Clinical data were abstracted from medical records including date at diagnosis, date at surgery, tumor stage, grade, histological estrogen receptor (ER), progesterone receptor (PR) and human epidermal growth factor (HER2) status, second cancers, breast-cancer recurrence, parity, history of breast feeding, age at menarche, and cause of death, if applicable. Six of the 140 breast-cancer patients had two primary contralateral breast cancers with tissue available for study for a total of 146 tumors.

### DNA and RNA sequencing

#### DNA extraction

Germline DNA was extracted from peripheral blood cells or from formalin-fixed paraffin-embedded (FFPE) normal breast tissue adjacent to tumor tissue from surgery. Peripheral blood cell DNA was extracted using a standard phenol chloroform method. For FFPE tissue, DNA and RNA were extracted from ten 30-μm sections from each tumor using the QIAmp DNA FFPE Tissue Kit (Qiagen) and miRNeasy Kit (Qiagen) according to manufacturer’s instructions. DNA was quantified with the Quant-iT PicoGreen dsDNA Assay Kit (Thermo Fisher Scientific, MA). After extraction and quantification, DNA was sent to The National Cancer Institute (NCI) Cancer Genomics Research Laboratory (CGR) for WES. For RNA sequencing, 500 ng total RNA was sent to the COH Integrative Genomics Core (IGC).

#### DNA library construction, hybridization, and massively parallel sequencing

Library production and sequencing for 146 tumors and 140 matching normal samples was performed at CGR. The KAPA HyperPlus Kit (Kapa Biosystems, Inc., Wilmington, MA) was used to generate libraries from 300ng DNA according to the KAPA-provided protocol. Libraries were pooled and sequence capture was performed with NimbleGen’s SeqCap EZ exome v3 (Roche NimbleGen, Inc., Madison, WI, USA), according to the manufacturer’s protocol. The resulting post-capture enriched multiplexed sequencing libraries were used in cluster formation on an Illumina cBOT (Illumina, San Diego, CA, USA) and paired-end sequencing was performed using an Illumina HiSeq 4000 following Illumina-provided protocols for 2 × 100 bp paired-end sequencing to an average-fold coverage of 80X for the tumors and 30X for the germline samples. Paired□ end reads from each sample were aligned to human reference genome (hg19) using Novoalign (v3.00.05), and the aligned binary format sequence (BAM) files were sorted and indexed using SAMtools (1, 2). The sorted and indexed BAMs were processed by Picard (v1.126, https://broadinstitute.github.io/picard/) to remove duplicate sequencing reads. Local realignment around suspected sites of indels was performed using Genome Analysis Toolkit (GATK) IndelRealigner (v3.3-0-g37228af). These mapped sequence reads were then base-recalibrated before being used for somatic mutation calling by MuTect2 in GATK (v4.0.11.0).

#### RNA-seq

In the COH IGC, sequencing libraries were prepared with Kapa RNA HyperPrep kit with RiboErase (Roche) and sequenced on a HiSeq 2500 (Illumina) with 40 million reads per sample. The RNA-seq reads were aligned to hg19 genome assembly using Tophat2 (v2.0.8) with default settings. The gene-expression levels were counted by obtaining raw counts with HTSeq (v0.6.1p1) against Ensembl v86 annotation. The counts data were normalized using the trimmed mean of M values (TMM) method implemented in R package edgeR^20^. Log2-transformed counts were used to assign PAM50 subtypes based on the subgroup-specific gene centering method developed by Zhao, *et al*.^21^. We estimated Z-scores based on the corrected median absolute deviation (MAD) implemented by the *robStandardize* R function in the robustHD R package and defined expression outliers as gene-sample data points with robust Z-scores greater than three. Raw counts of RNA-seq data for 1,189 TCGA samples (including both tumor and matched normal samples) were downloaded from the Genomic Data Commons (GDC) using the GDCRNATools^22^ R package. RNA-seq data for H/L tumor samples and TCGA samples were processed and analyzed separately.

### Data analysis

#### Germline variant calling

Germline variant calling from the BAM files was performed in the COH IGC using GATK HaplotypeCaller (https://software.broadinstitute.org/gatk). Variants with a call quality less than 20, read depth less than 10, or allele fraction ratio less than 20% were removed. Variants in variant call format files were evaluated for pathogenicity using Ingenuity Variant Analysis (IVA) version 4 (Qiagen Inc, Alameda, CA) and American College of Medical Genetics and Genomics (ACMGG) guidelines were applied using the IVA ACMGG calling algorithm^23^. Pathogenic or likely pathogenic variants were individually evaluated by the research team using the available literature and ClinVar to make a final determination^24^.

#### Genetic ancestry analysis

We performed genetic ancestry estimation for each of the 140 women using the germline whole-exome sequencing data. We used 90 European (1000 Genomes), 90 African (1000 Genomes), 90 East-Asian (1000 Genomes) and 71 IA ancestry^25^ reference samples. We identified the SNPs that overlap all data sets (N=9,935). We combined all SNPs and dropped SNPs that did not match based on reference and alternate alleles. To estimate the ancestry for each sample, we used ADMIXTURE 1.3.0 setting the K parameter to 4 and running the unsupervised algorithm^26^. In addition, we used principal components analysis, calculated using PLINK 1.9^27^ as a complementary method to assess ancestry.

#### Somatic variant calling

We identified somatic single nucleotide variants (SNV) using MuTect2 in GATK4 (v4.0.11.0) suite with default parameters^28^ and indels using GATK Indelocator. Using the SNV and indel filtering method described in Pereira et al.^3^, we focused on frameshift, non-synonymous, canonical splicing site, and stop gain mutations. Briefly, somatic mutations were manually curated and considered true positives in a sample if the mutation was observed in >10% of reads or with a frequency of 5-10% if in frequently mutated breast-cancer genes or seen in COSMIC database^29^. Because the tumors include both tumor and normal stromal cells, it is expected that the proportion of reads will have less than the expected 50% if 100% tumor. Mutations in <5% of reads, in segmental duplication regions, or indels that overlapped homopolymer stretches of six or more bases were considered false positives. We did visual checking using the Integrative Genomics Viewer (IGV) to assess the quality of all somatic mutations. We performed Sanger sequencing on a subset of samples to confirm specific mutations in *AKT1, BARD1, MAP3K1*, and *MET*. Using the filtered and annotated somatic mutations, we performed a somatic-mutation significance analysis via MutSigCV^30^ (version 1.3.5) on Genepattern (https://www.genepattern.org/modules/docs/MutSigCV). Genes with false discovery rate (FDR) q < 0.05 are considered to be significantly mutated genes.

We compared the significant somatic mutations in our analysis with the mutations from the Romero-Cordoba dataset^19^. Using the publicly available somatic-mutation data from the Romero-Cordoba study of the Mexican patients, we combined our somatic-mutation data and performed a MutSigCV analysis to identify the common significant genes. Similarly, to investigate if these significantly mutated genes were associated with ancestry, we performed the same analysis on breast tumors from Whites in TCGA. Using 2% as the mutation frequency threshold, we performed Fisher’s exact test for each frequently mutated gene for comparison.

#### Copy-number analysis using FACETS

We used FACETS implemented in R package facets version 0.6.1^31^ to calculate CNAs. The counts of reads with the reference (ref) allele, alternate (alt) allele, errors (neither ref nor alt), and deletions at a specific genomic position were generated using BAM files from the 146 matched tumor-normal sample pairs using the application snp-pileup in the facets package. The segmentation of each tumor sample was then estimated with the critical value (cval) 150.

The segmentation files generated by facets served as input files for the GISTIC2.0^32^ on the GenePattern server (https://genepattern.broadinstitute.org/gp) to identify significant SCNAs using a q-value cutoff < 0.05. A gene was considered as copy number altered with GISTIC2-thresholded scores of −2 (deep loss), −1 (shallow loss), 1 (low-level gain) and 2 (high-level gain). The GISTIC2 copy-number results and clinical data for 816 TCGA tumor samples were downloaded from the cBioPortal database^33^ (https://www.cbioportal.org). Expression outliers (defined by Z-scores greater than 3.0) were considered as driven by copy-number changes if greater than 90% expression outliers in a gene had a GISTIC2-thresholded copy-number score of 2 (high-level gain) or 1 (low-level gain). Fisher’s exact test was used to identify genes with frequency difference in expression outliers, driven by copy-number alterations, between 146 tumor samples from H/L and 452 TCGA Whites (determined as having > 95% European ancestry as described below).

#### Mutation-signature analysis

Using the previously called SNVs, we performed a mutational signature analysis via the MutationalPatterns R package^34^. Hg19 was used as the reference genome. SNVs were parsed and classified into six mutation patterns (C>T, T>A, C>G, T>C, C>A and T>G) and 96 trinucleotide changes. Then a non-negative matrix factorization algorithm was implemented to extract mutation signatures. And we compared the similarities of these mutation signatures with the COSMIC mutation signatures and each mutation signature could be treated as a linear combination of the 30 COSMIC mutation signatures. The 30 COSMIC mutation signature percentage contribution was then computed for each tumor and a contribution heatmap was generated. Within these tumor samples, we performed a signature contribution comparison using the two-sided Wilcoxon rank-sum tests among the five tumor subtypes (Luminal A, luminal B, basal-like, HER2-enriched and normal-like).

We also compared the mutation-signature analysis with the breast tumors in the Romero-Cordoba dataset and the breast tumors from Whites in TCGA SNV dataset. For the significant COSMIC mutation signatures identified in our dataset, we performed two-sided Wilcoxon rank-sum tests among the three datasets to test if the signature was enriched in Mexican patients.

## RESULTS

### Clinical/demographic data and germline pathogenic variants

Characteristics of the 140 participants are shown in Table 1. The mean age at diagnosis was 48.7 years with a range from ages 31 to 75 years. Nearly all of the 140 H/L were of mixed European (Eur) and IA ancestry. The mean ancestry composition was 50.6% Eur, 40.8% IA, 5.9% African, and 2.7% Asian although the range of ancestry proportion varied widely from <1% to 96% IA at the extremes (Figure 1). As shown in the Principal Component Analysis (PCA) plots in Figure 1A, H/L samples are not well-represented in TCGA project. For the six individuals with two primary tumors (in the contralateral breasts), the tumors were considered independent tumors (Supplemental Table 1) which was borne out by different somatic-mutation profiles. The majority of the women were diagnosed with Stage I (44%) or II (43%) tumors (Table 1). There were 22 recurrences and 10 deaths during the time of follow-up. Of the 146 tumors, 83% were ER-positive, 72% were PR-positive, and 17% were HER-2 positive and these proportions were similar to White women in TCGA ^1^. Germline pathogenic variants in breast cancer predisposition genes were identified in six participants including one *BRCA1* exon 9-12 deletion, four *CHEK2* L236P, and one *NF1* Y408X variants of which the *BRCA1* and *CHEK2* variants are of Indigenous-American ancestry^35^.

**Table 1.** Patient and tumor characteristics of 140 H/L breast-cancer cases and their 146 breast tumors

| Patient characteristics | Mean | Range | Median |  |
| --- | --- | --- | --- | --- |
| Age at diagnosis (years) | 48.7 | 31-75 | 48 |  |
| Breastfeeding (months) | 7.2 | 0-84 | 2 |  |
| Parity (number children) | 2.3 | 0-8 | 2 |  |
| Age at menarche (years) | 12.6 | 9-18 | 12 |  |
| **Tumor characteristics** | Positive | Negative | Unknown | Equivocal |
| Estrogen receptor | 120 (82%) | 25 (17%) | 1 (0.7%) |  |
| Progesterone receptor | 104 (72%) | 41 (28%) | 1 (0.7%) |  |
| HER2 | 25 (17%) | 116 (80%) | 1 (0.7%) | 4 (3%) |
| Stage at diagnosis | I | II | III | IV |
|  | 63 (44%) | 63 (43%) | 17 (12%) | 3(2%) |

**Figure 1.**
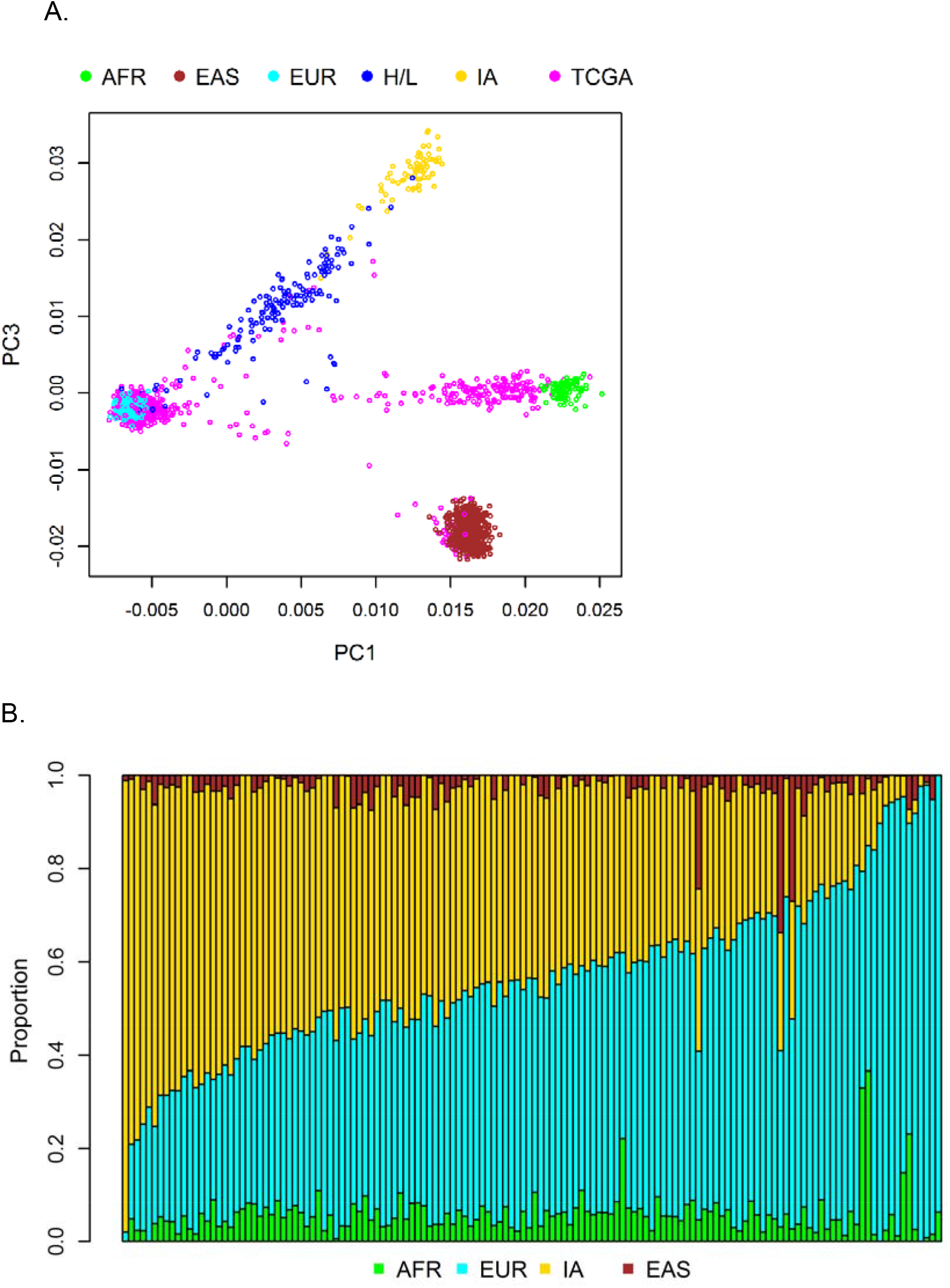
Ancestry of the cohort. Results of principal components analysis comparing the values for samples on principal component (PC) 1 (x-axis) and PC3 (y-axis) (A). Each dot represents the results from one individual: Hispanic/Latina (H/L) (dark blue); TCGA (pink); and reference populations including African (AFR), Yoruban individuals from Nigeria from HapMap (light green); East Asians (EAS), Han Chinese from HapMap (brown); European American (EUR), CEBP from HapMap (light blue); and Indigenous American (IA) (yellow) from Mexico. PC2 (not shown) captures individuals of Asian and Indigenous-American ancestry. (B). Results from ADMIXTURE analysis. Each vertical bar represents estimate of ancestry from one individual. Ancestry is assigned for each individual as a fraction of either African (green), Asian (brown), European (light blue) or Indigenous American (yellow) ancestry.

### Somatic mutations

We observed a total of 4510 true somatic mutations in 3391 genes in the 146 primary breast tumors (Supplemental Table 2). The number of mutations per individual varied from 2 to 225. Using MutSigCV, we found that mutations in *PIK3CA, TP53, GATA3, MAP3K1, CDH1, CBFB, PTEN*, and *RUNX1* were significant (FDR < 0.05) cancer driver mutations. To identify additional, potentially significantly mutated genes in H/L, we merged the mutation data from our cohort with a previously published study of Mexican breast-cancer patients (N = 135)^19^. Within the aggregated mutation data of this combined cohort (N = 281), we re-ran MutSigCV and identified one more significantly mutated gene, *AKT1*, which only occurred twice in our 146 primary breast tumors. Using the statistically significantly mutated genes obtained from the aggregated cohort, we visualized the mutational profiles within our cohort (Figure 2a) and the variant locations for *PIK3CA* and *GATA3* (Figure 2b). For *MAP3K1* and *RUNX1*, at least one tumor harbored multiple mutations in the same gene. Furthermore, in *GATA3*, eight tumors had the identical splice mutation (NM_002051.2:exon5:r.spl;NM_001002295.1:exon5:r.spl) that affected expression (data not shown). Other genes of interest that did not meet the significance threshold (FDR < 0.05) but which have been identified as significant in prior studies and were mutated in our dataset included *MLL3* (aka *KMTC2*) (6%), *PTPRD* (3%), *MAP2K4* (2%), *PIK3R1* (2%), *NF1* (1%), *RB1* (1%), *TBX3* (1%), *FOXA1* (1%), *PADI4* (1%), *CDKN1B* (1%), *CTCF* (1%), and *NCOR1* (1%). In addition, we found mutations in *MET* (4.1%) which is not generally considered a breast-cancer gene but is a known driver in other cancer types^36^.

**Table 2:**
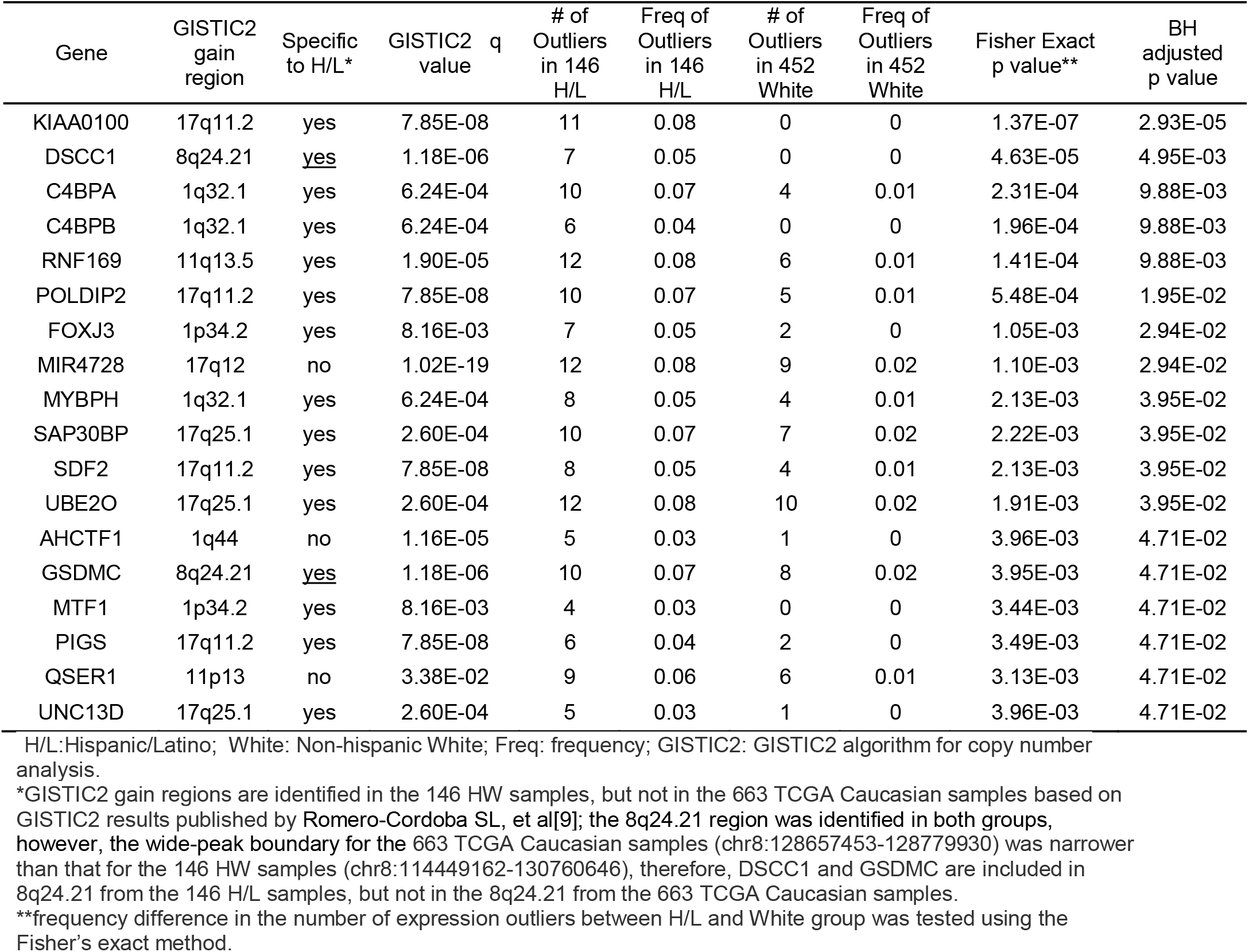
Frequency difference in expression outliers driven by copy number gain between 146 tumors from H/L and 452 tumors from TCGA White

**Figure 2.**
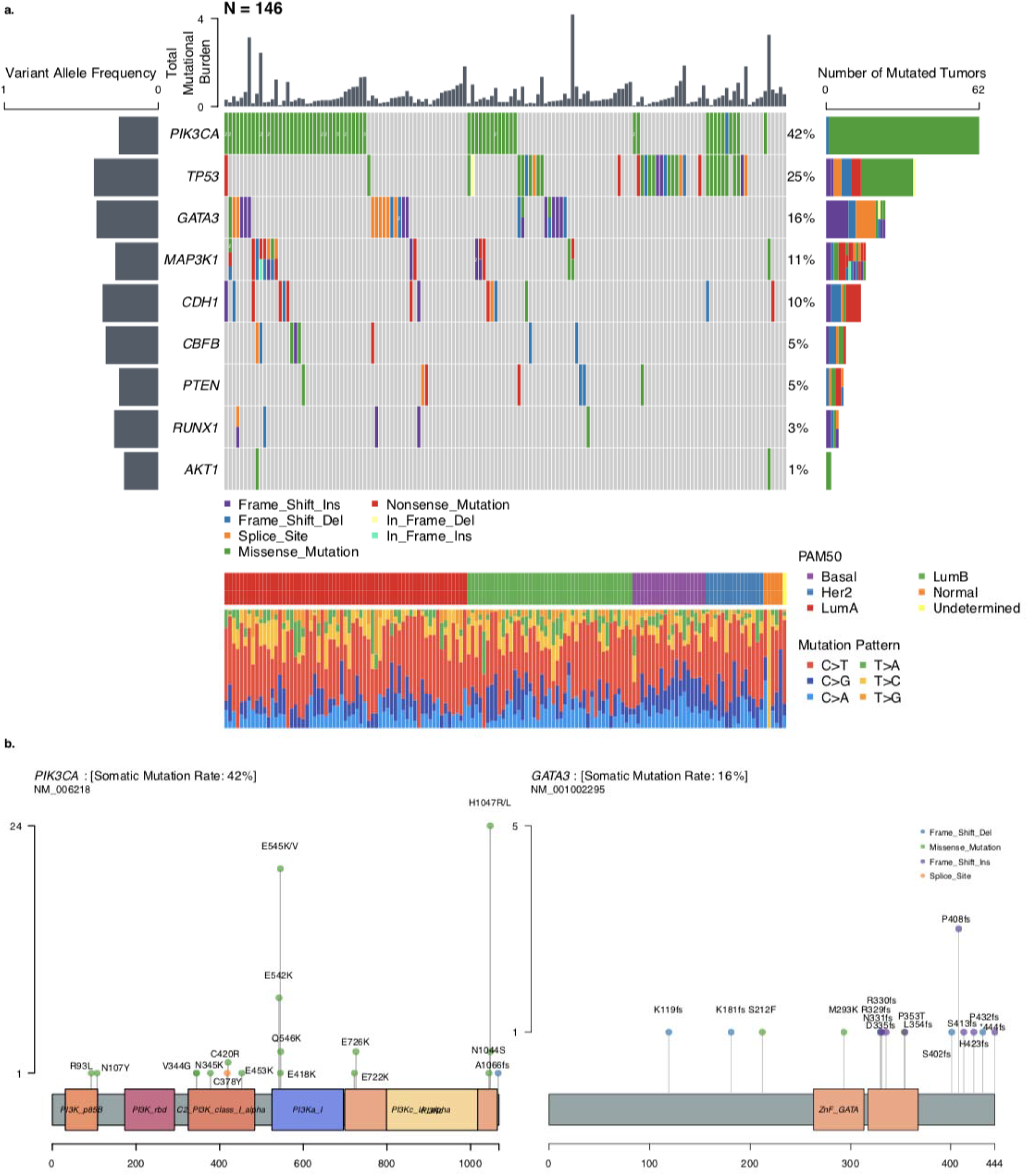
Tumor mutational burdens and somatic-mutational profiles. a. Mutation plot of nine significantly mutated genes in the 146 tumors. Different mutation classifications are color-coded Numbers are shown where multiple mutations of the same classification were detected. Total mutational burden for each tumor is shown as a bar chart on top. The mean variant allelic frequency is shown for each gene on the left. PAM50 subtype and mutation pattern for each tumor are shown at the bottom. b. Lollipop plot of *PIK3CA* and *GATA3* mutations within the 146 tumors. Mutation classifications are color coded and amino-acid changes are specified for each mutation.

The frequency of mutations in genes known to be significantly mutated in breast cancer, including *PIK3CA, MAP3K1, GATA3, CBFB, and MLL3/KMT2C*, were not significantly different in tumors from H/L compared to tumors from White women in TCGA (FDR q > 0.05, Supplemental Table 3). Similar to tumors from Whites, *PIK3CA* and *TP53* were the most commonly mutated genes. We identified *AKT1* mutations in 2 of 146 tumors (1.4%), including the E17K hotspot mutation which was found to be mutated in 8% of patients among Mexican women^19^. After correction for multiple hypothesis testing, we found no somatic mutations significantly associated with genetic ancestry.

### Mutational signature analysis

To investigate the mutational processes in H/L breast-cancer tumors and the association between PAM50 subtypes and mutational patterns, we adopted the non-negative matrix factorization approach as proposed by Alexandrov et al.^37^ for mutational signature analysis of tumors. Signature calling revealed five major contributing signatures in the 146 tumors corresponding to the COSMIC signatures 1,2,3,13 and 16 (Figure 3a; Supplemental Table 4). Signature 1 was detected in all 146 tumors. The contribution of COSMIC signature 1 was greater in luminal A and B subtypes than HER2 and basal subtypes (p < 0.05, two-sided Wilcoxon rank sum test) (Figure 3b). Signatures 2 and 13, attributed to activity of the AID/APOBEC family of cytidine deaminases, were found in tandem in 16% (n = 23) of the tumors and were more common in tumors with HER2 subtype compared to luminal A and B subtypes (Figure 3b). We found that 13 tumors were homozygous and 29 tumors were heterozygous for a common 29.5kbp germline deletion spanning most of *APOBEC3B*. Tumors with the deletion had a higher proportion of COSMIC signatures 2 (p = 0.0005, Wilcoxon rank sum test) and 13 (p = 0.0008, Wilcoxon rank sum test). Signature 3, attributed to defects of homologous recombination double-stranded DNA break-repair, was found significantly more often in basal subtypes than the other PAM50 subtypes (p < 0.05, two-sided Wilcoxon rank sum test) including the tumor with the germline *BRCA1* exon 9-12 deletion. We observed a group of tumors (N=40, 27.4%) with more than 5% COSMIC signature 16 contributions. Since this was not previously reported in other breast tumor studies, we re-examined other datasets, using the same analytic pipeline used herein. We found that signature 16 was present in 20 (19.6%) tumors in a previous study of Mexican breast-cancer patients^19^ which was not significantly different than the proportion in our dataset (p = 0.18, Fisher’s exact test). The proportion in tumors from TCGA White women (N=75; 8.9%) was significantly lower than in our dataset (p < 0.001, Fisher’s exact test) (Figure 3c) and in the Romero-Cordoba et al. dataset (p < 0.0001, Fisher’s exact test). The percentage of this signature was significantly higher in luminal A and B subtypes compared to HER2 and basal tumors (p < 0.05, two-sided Wilcoxon rank sum test) (Figure 3b).

**Figure 3.**
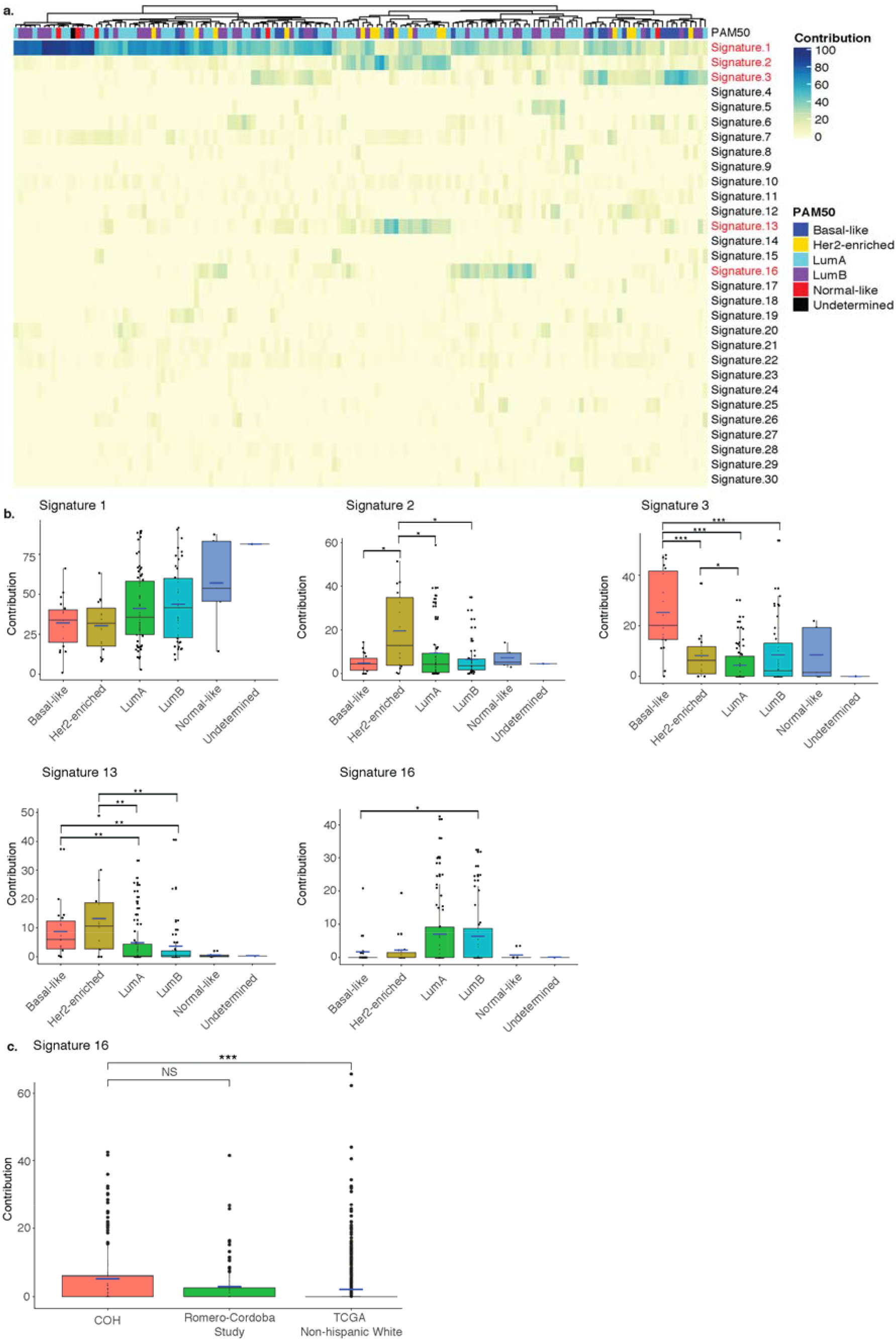
Mutational Signatures. a. Unsupervised clustered heatmap of contributions from each mutational signatures for the 146 tumors. Significant signatures are highlighted in red. PAM50 subtype for each tumor is shown on top of the heatmap. b. Box-plots comparison of the contributions of the five significant mutational signatures (Signature 1, 2, 3, 13, 16) across the PAM50 subtypes. Statistical significance levels are indicated within the box plots (*: p < 0.05; **: p < 0.01; ***: p < 0.001, Wilcoxon Rank-sum test). c. Box plot of Signature 16 contributions in the 146 tumors from the Hispanic-Mexican cohort (COH), Romero-Cordoba Study and the Non-Hispanic White tumors in the TCGA dataset. Statistical significance levels are indicated within the box plot (NS: not significant, p > 0.05; *: p < 0.05; **: p < 0.01; ***: p < 0.001, Wilcoxon Rank-sum test).

### Somatic CNAs (SCNAs)

Using GISTIC2, we identified chromosome arm-level SCNAs that were significantly (q < 0.05) amplified at 1q, 8q, 6p, 1p, 6q, 16p, 20q, 8p, 12q and deleted at 22q, 16p, 17p, 8p (Supplemental Table 5). In addition to these broad SCNAs, we identified significantly (q < 0.05) amplified or deleted focal regions including 29 peak regions of amplification and 48 regions of deletion (Figure 4A). Seven recurrently amplified regions contain common oncogenes (*FGFR1, MYC, CCND1, MDM2, IGF1R, ERBB2*, and *ZFP217*); one recurrently deleted region contains *TP53* (Figure 4A). By integrative analysis of RNA-seq gene-expression data and copy-number data, we observed that greater than 90% of expression outliers (defined by robust Z-score greater than 3.0) in *ERBB2, FGFR1, IGF1R*, and *MDM2* were associated with copy-number gain (Figure 4B). Therefore, we sought to identify expression outliers from 1,121 genes contained in the 29 copy-number amplification peak regions for the 146 H/L breast tumor samples and 452 White TCGA breast tumor samples. Of 1,121 genes in the 29 regions, over 90% of expression outliers were associated with copy-number gain in 214 genes, including 88 genes from the 146 H/L samples, 62 genes from the 452 TCGA White samples, and 64 genes from both sample groups (Supplemental Table 6). Eighteen of 214 genes had significant (FDR < 0.05) difference in frequency of expression outliers copy number between the 146 H/L and 452 TCGA White tumor samples (Table 2 and the top 18 rows in Supplemental Table 6). Expression outliers from those genes were more prevalent in the 146 H/L than in the 452 White tumors because we focused on the 29 copy-number regions (Figure 4A) found in H/L (Table 2; Supplemental Table 6).

**Figure 4:**
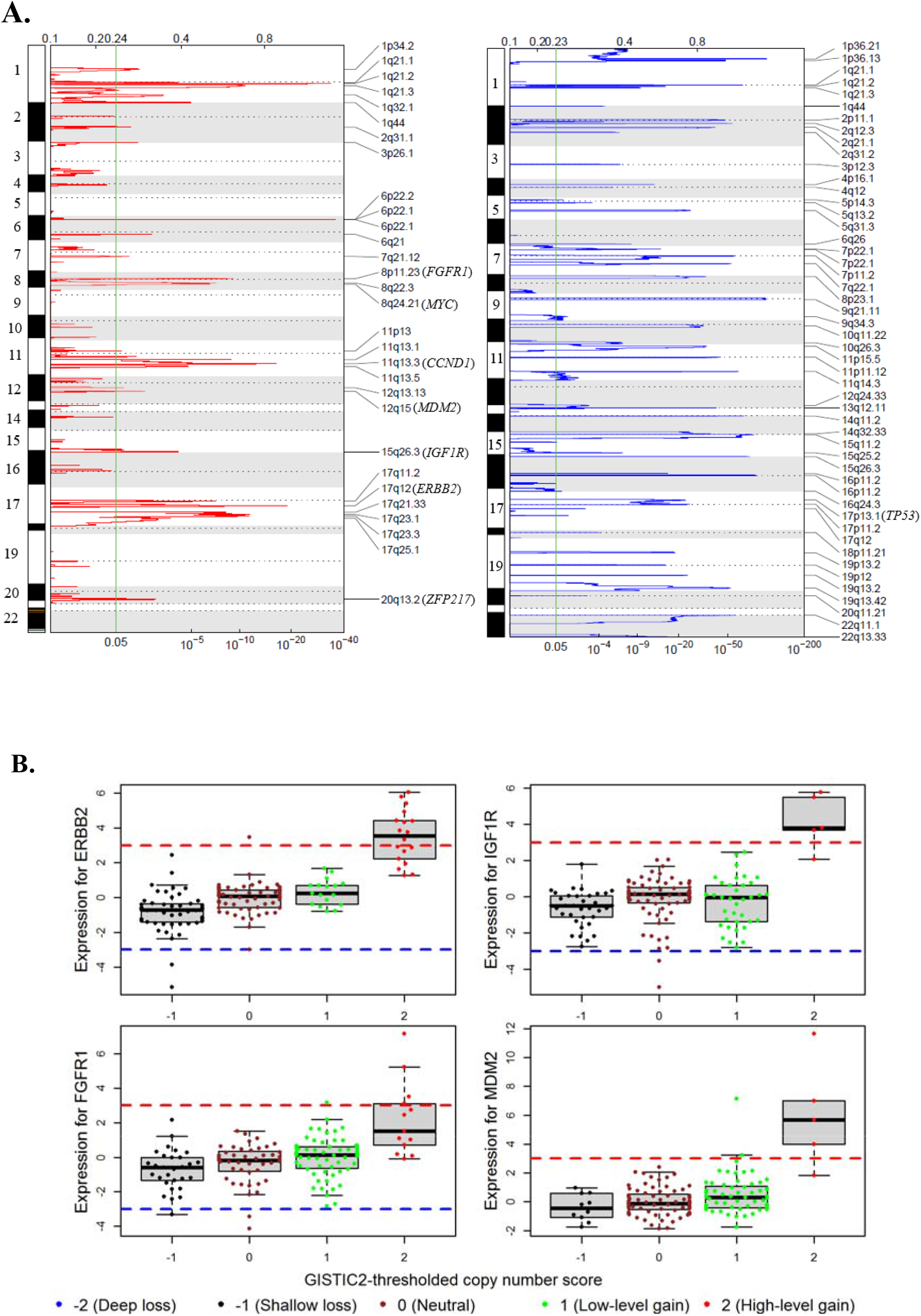
Copy-number alterations. a. Genomic regions of significant copy-number gain (left) and loss (right) identified by GISTIC2. Common oncogenes and tumor suppressor genes are in parentheses next to the corresponding cytobands. The green vertical line marks the GISTIC2 q value of 0.05 (bottom x-axis). b. Outlying gene expression and copy-number gain in four genes in 146 H/L breast tumor samples. Gene-expression values on the y-axis are Z-scores estimated by robust standardization; the red dash line of Z-score = 3 and blue dash line of Z-score = −3 are cutoff values for outliers of over-expression and under-expression, respectively.

Using this combined copy-number and gene-expression analysis approach, we identified *KIAA0100*, also known as Breast Cancer Overexpressed Gene 1 (*BCOX1*), as the top gene that was systematically different between Whites (TCGA) and our H/L cohort. (Figure 5A). Since this gene is within ∼11 megabases of *ERBB2* on chromosome 17q, we investigated whether it was part of the ERBB2 GISTIC amplification peak. We observed that the peaks for copy-number amplifications (Figure 5B) were distinct for *KIAA0100* and *ERBB2* are at 17q11.2 and 17q12.

**Figure 5.**
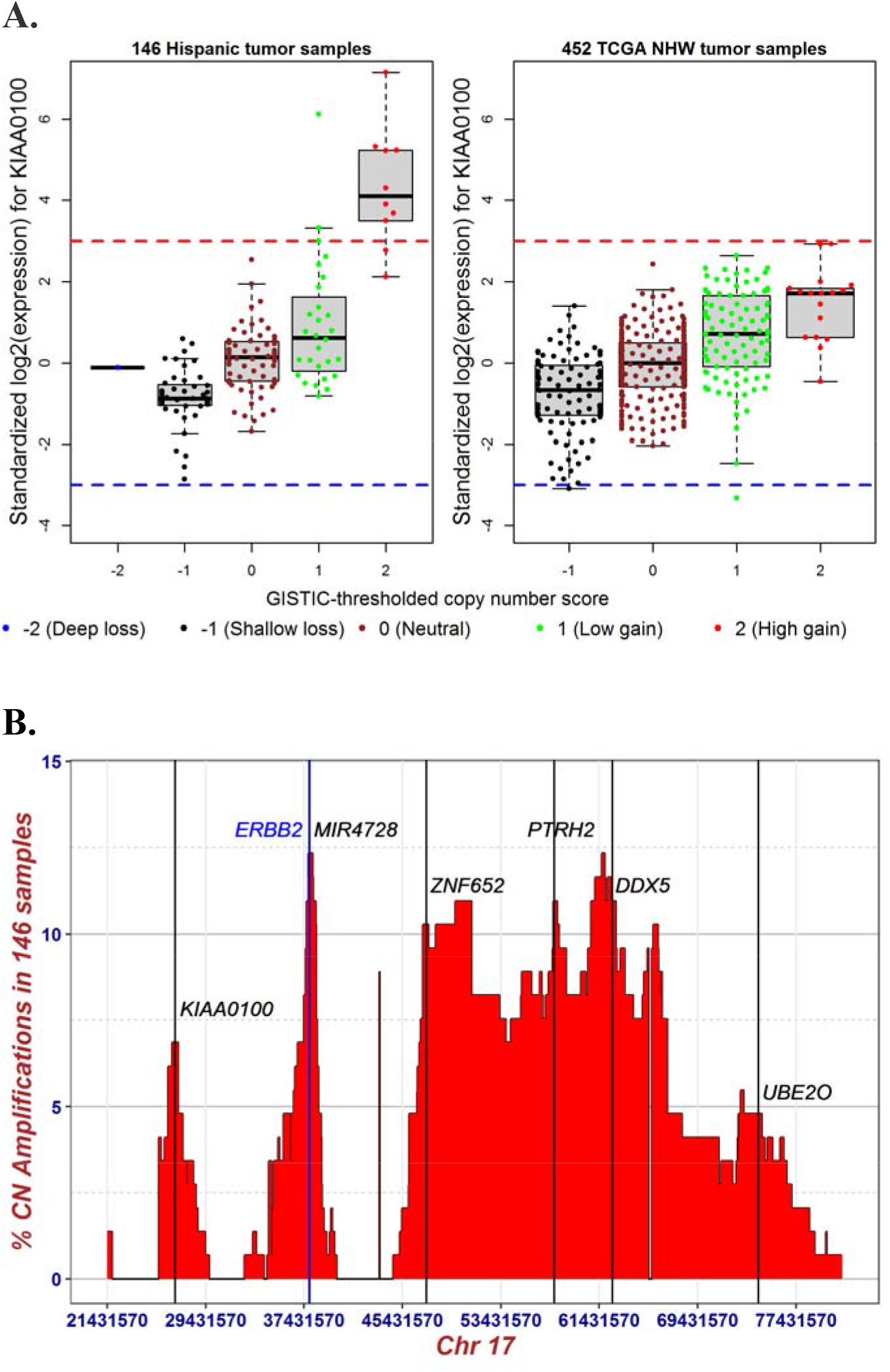
Expression outliers and copy-number gain in KIAA0100. a. Distribution of gene expression and GISTIC2-thresholded copy-number scores in *KIAA0100* for 146 breast tumor samples from Hispanic whites and 452 breast tumor samples from TCGA Non-Hispanic whites (A). The y-axis is standardized gene-expression values (Z-scores) estimated robustly based on the corrected median absolute deviation (MAD). Red and blue dashed lines represent Z-score of 3 and −3, respectively. b. Distribution of proportion of high-level copy-number gain for 950 genes spanning the 6 amplified regions of 17q11.2, 17q12, 17q21.33, 17q23.1, 17q23.3, and 17q25.1. Y-axis is the percentage of the 146 Hispanic samples with GISTIC2-thresholded copy-number score of 2; x-axis is genomic boundaries (Chr17: 21431570 – 81188573, hg19) for the six significantly amplified regions determined by GISTIC2. The vertical lines mark the genomic locations of *KIAA0100* (*BCOX1*, 17q11.2) at Chr17:26941457 – 26972177, *ERBB2* (17q12) at Chr17: 37844336 – 37873910, *MIR4728* (microRNA 4728, 17q12) at Chr17: 37882747 – 37882814, *ZNF652* (17q21.33) at Chr17: 47366567 – 47439476, *PTRH2* (17q23.1) at Chr17: 57774666 – 57784959, *DDX5* (17q23.3) at Chr17: 62494371 – 62503156, and *UBE2O* (17q25.1) at Chr17: 74385612 – 74449288.

## Discussion

We analyzed tumor-germline sequencing data combined with RNA-seq data from 146 tumors from 140 self-identified H/L recruited from a single center in the Los Angeles region. As expected, the majority were of mixed European and IA ancestries. Since TCGA has extremely limited samples of breast cancer from H/L and particularly of H/L of mixed IA ancestry, our report fills a critical gap in the landscape of somatic mutations and copy-number alterations in this increasing US population. Together, our analyses and the recent paper focused on Mexican women living in Mexico ^19^ substantially enhance the data in the public domain for women of H/L heritage.

The most commonly mutated gene in our population was *PIK3CA* which is the most commonly mutated gene in TCGA White samples. For women with advanced ER+/HER2-breast cancers, alpelisib is a currently approved therapy, and our results suggest that this therapy should be useful in a large fraction of H/L women. The Romero-Cordoba et. al. study identified a high frequency (8%) of the E17K activating *AKT1* mutation indicating such women may benefit from AKT inhibitors. We only identified two tumors with mutations in *AKT1* and only one with the E17K mutation. The difference between our results and those of Romero-Cordoba may be due to chance, differences in selection criteria between the two cohorts, and/or differences in environmental exposures between the two cohorts. Since the ancestry of our population is similar, it is unlikely that the differences we observed are due to germline-genetic differences between the two cohorts.

We performed analyses of the somatic-mutational signatures and compared them to the TCGA dataset. Our analysis identified COSMIC signature 16 (contribution > 5%) in a significant fraction of tumors (27.4%) in our dataset with similar rates in the data from Romero-Cordoba et al. who analyzed breast tumors from Mexican patients. Because Romero-Cordoba et al. used a contribution cutoff in their mutation-signature-analysis pipeline, they did not report this signature. However, in our analysis, we implemented the non-negative matrix factorization algorithm and no contribution cutoff was applied such that signature 16 was observed. There were significantly lower rates of this signature in TCGA White women (p< 0.001). We do not believe our finding is a technical artifact from FFPE because this signature was found in frozen tissue in the Romero-Cordoba et al data. No known genetic or environmental exposures that predispose to this signature have been reported and prior studies have not found this mutational signature in breast cancer, although it has been reported to be common in liver cancers^37^.

Other COSMIC signatures were the same as those previously reported in TCGA. We found signatures 2 and 13 associated with APOBEC loss as a relatively common finding, associated with HER2-amplified tumors and specifically with the germline APOBEC copy-number variant similar to previous reports^38^. The APOBEC*3B* common 29.5-kbp germline deletion results in the fusion of *APOBEC3A* and the 3’UTR of *APOBEC3B*^*39*^. This fusion generates a more stable *APOBEC3A* mRNA, resulting in increased expression of *APOBEC3A*, higher overall mutation burden, and a higher odds ratio of developing breast cancer^40, 41^. We also found Signature 3, associated with defects in homologous recombination repair as a common signature, which is over-represented in basal-like tumors as previously reported^37, 42^.

Our copy-number analyses identified copy-number gains, i.e., 1q, 8q, 17q which are common in breast cancer in other populations^1, 2^. We also identified several known CNAs which were recurrently gained in our dataset. In combined analysis of copy-number alterations and gene expression, we identified *KIAA0100* (*BCOX1*) as a recurrently amplified region with high gene expression which was more common in tumors from H/L than tumors from White women in TCGA. KIAA0100 was originally identified in a screen for genes that were more frequently found in breast tumor than in normal breast tissue^43^ and increased expression was associated with poor prognosis^43, 44^. Knock-down of *KIAA0100* by siRNA in the breast-cancer cell line MDA-MB-231 reduced cell aggregation, reattachment, cell metastasis and invasion^45^. Thus, KIAA0100 may be of interest for further study in understanding the biology of tumors in H/L and stratifying women for risk of recurrence.

Our study has several limitations. We included only women who did not have neoadjuvant therapy prior to surgical resection. We chose this subset of women to avoid effects possibly induced by neoadjuvant chemotherapy such as new mutations and/or selection for resistant subclones. However, because neoadjuvant therapy is more likely to be given to patients with large tumors and/or tumors with poor prognosis^46^, tumors included in our study may have some differences in comparison with prior studies due to these selection criteria. For example, because most triple-negative breast tumors are first treated with neoadjuvant therapy, the proportion of triple-negative tumors in our study was lower than previously reported^47^. Our analysis of tumor copy-number alterations was based on WES data. Although WES and other forms of targeted sequencing are used for CNA analysis, it makes it difficult to conduct one-to-one comparisons to array-based or whole genome sequencing-based analyses. Therefore, we limited our analyses to copy-number events that also demonstrated gene-expression differences across populations. Finally, although our study substantially increases the number of tumors analyzed by WES in H/L, the overall numbers are still substantially lower than in White women. In particular, we are likely underpowered to discover low frequency, ethnic and/or ancestry-specific drivers that may be unique to this population. There also were too few recurrences and deaths for statistical analyses.

In summary, we conducted a comprehensive characterization of somatic mutations, CNAs, and gene expression in 146 breast tumors from 140 H/L from Los Angeles County, California. We found that COSMIC signature 16 was more common in our dataset and a recently published dataset of Mexican women living in Mexico, suggesting that this signature may be important in self-reported H/L/Hispanic women and potentially useful to understand differences at diagnosis and for outcome. Finally, our combined CNA and gene-expression analysis suggested that KIAA0100 may be a possible driver of breast-cancer aggressiveness in a subset of our sample. These results should be useful to understanding the biology and guiding therapy for breast cancer among H/L.

## Acknowledgments

This work was funded by the National Cancer Institute (R01CA184585, K24CA169004), the National Institute on Minority Health and Health Disparities Division of Intramural Research, and the California Initiative to Advance Precision Medicine (OPR18111). Research reported in this publication included work performed in the City of Hope Integrative Genomics Core and the Pathology Core supported by the National Cancer Institute of the National Institutes of Health under grant number P30CA033572. The content and views are solely the responsibility of the authors and should not be construed to represent the views of the National Institutes of Health. SLN and this research were partially funded by the Morris and Horowitz Families Professorship. CDA is supported by the National Heart, Lung, and Blood Institute (NHLBI T32HL007118) through the training Program in Molecular and Integrative Physiological Sciences at the Harvard T.H. Chan School of Public Health. LF is supported by R01CA204797. JNW was supported by NIH RC4 CA153828; Breast Cancer Research Foundation (#20-172), and American Society of Clinical Oncology Conquer Cancer® Research Professorship in Breast Cancer Disparities.

## Conflicts of interest

JNW is a speaker for the Bureau for AstraZeneca, and an employee at Natera. No other conflicts of interest from authors

Data are being deposited in dbGAP and will be made available at the time of acceptance.

## References

1. Cancer Genome Atlas N. Comprehensive molecular portraits of human breast tumours. Nature. 2012;490(7418):61–70. Epub 20120923. doi: 10.1038/nature11412. PubMed PMID: 23000897; PMCID: PMC3465532.

2. Curtis C, Shah SP, Chin SF, Turashvili G, Rueda OM, Dunning MJ, Speed D, Lynch AG, Samarajiwa S, Yuan Y, Graf S, Ha G, Haffari G, Bashashati A, Russell R, McKinney S, Group M, Langerod A, Green A, Provenzano E, Wishart G, Pinder S, Watson P, Markowetz F, Murphy L, Ellis I, Purushotham A, Borresen-Dale AL, Brenton JD, Tavare S, Caldas C, Aparicio S. The genomic and transcriptomic architecture of 2,000 breast tumours reveals novel subgroups. Nature. 2012;486(7403):346–52. Epub 20120418. doi: 10.1038/nature10983. PubMed PMID: 22522925; PMCID: PMC3440846.

3. Pereira B, Chin SF, Rueda OM, Vollan HK, Provenzano E, Bardwell HA, Pugh M, Jones L, Russell R, Sammut SJ, Tsui DW, Liu B, Dawson SJ, Abraham J, Northen H, Peden JF, Mukherjee A, Turashvili G, Green AR, McKinney S, Oloumi A, Shah S, Rosenfeld N, Murphy L, Bentley DR, Ellis IO, Purushotham A, Pinder SE, Borresen-Dale AL, Earl HM, Pharoah PD, Ross MT, Aparicio S, Caldas C. The somatic mutation profiles of 2,433 breast cancers refines their genomic and transcriptomic landscapes. Nat Commun. 2016;7:11479. Epub 20160510. doi: 10.1038/ncomms11479. PubMed PMID: 27161491; PMCID: PMC4866047.

4. Andre F, Ciruelos E, Rubovszky G, Campone M, Loibl S, Rugo HS, Iwata H, Conte P, Mayer IA, Kaufman B, Yamashita T, Lu YS, Inoue K, Takahashi M, Papai Z, Longin AS, Mills D, Wilke C, Hirawat S, Juric D, Group S-S. Alpelisib for PIK3CA-Mutated, Hormone Receptor-Positive Advanced Breast Cancer. N Engl J Med. 2019;380(20):1929–40. doi: 10.1056/NEJMoa1813904. PubMed PMID: 31091374.

5. Huang SF, Liu HP, Li LH, Ku YC, Fu YN, Tsai HY, Chen YT, Lin YF, Chang WC, Kuo HP, Wu YC, Chen YR, Tsai SF. High frequency of epidermal growth factor receptor mutations with complex patterns in non-small cell lung cancers related to gefitinib responsiveness in Taiwan. Clin Cancer Res. 2004;10(24):8195–203. doi: 10.1158/1078-0432.CCR-04-1245. PubMed PMID: 15623594.

6. Arrieta O, Cardona AF, Martin C, Mas-Lopez L, Corrales-Rodriguez L, Bramuglia G, Castillo-Fernandez O, Meyerson M, Amieva-Rivera E, Campos-Parra AD, Carranza H, Gomez de la Torre jc, Powazniak Y, Aldaco-Sarvide F, Vargas C, Trigo M, Magallanes-Maciel M, Otero J, Sanchez-Reyes R, Cuello M. Updated Frequency of EGFR and KRAS Mutations in NonSmall-Cell Lung Cancer in Latin America: The Latin-American Consortium for the Investigation of Lung Cancer (CLICaP). J Thorac Oncol. 2015;10(5):838–43. doi: 10.1097/JTO.0000000000000481. PubMed PMID: 25634006.

7. Carrot-Zhang J, Soca-Chafre G, Patterson N, Thorner AR, Nag A, Watson J, Genovese G, Rodriguez J, Gelbard MK, Corrales-Rodriguez L, Mitsuishi Y, Ha G, Campbell JD, Oxnard GR, Arrieta O, Cardona AF, Gusev A, Meyerson M. Genetic Ancestry Contributes to Somatic Mutations in Lung Cancers from Admixed Latin American Populations. Cancer Discov. 2021;11(3):591–8. Epub 20201202. doi: 10.1158/2159-8290.CD-20-1165. PubMed PMID: 33268447; PMCID: PMC7933062.

8. Li J, Xu C, Lee HJ, Ren S, Zi X, Zhang Z, Wang H, Yu Y, Yang C, Gao X, Hou J, Wang L, Yang B, Yang Q, Ye H, Zhou T, Lu X, Wang Y, Qu M, Yang Q, Zhang W, Shah NM, Pehrsson EC, Wang S, Wang Z, Jiang J, Zhu Y, Chen R, Chen H, Zhu F, Lian B, Li X, Zhang Y, Wang C, Wang Y, Xiao G, Jiang J, Yang Y, Liang C, Hou J, Han C, Chen M, Jiang N, Zhang D, Wu S, Yang J, Wang T, Chen Y, Cai J, Yang W, Xu J, Wang S, Gao X, Wang T, Sun Y. A genomic and epigenomic atlas of prostate cancer in Asian populations. Nature. 2020;580(7801):93–9. Epub 20200325. doi: 10.1038/s41586-020-2135-x. PubMed PMID: 32238934.

9. Yuan J, Hu Z, Mahal BA, Zhao SD, Kensler KH, Pi J, Hu X, Zhang Y, Wang Y, Jiang J, Li C, Zhong X, Montone KT, Guan G, Tanyi JL, Fan Y, Xu X, Morgan MA, Long M, Zhang Y, Zhang R, Sood AK, Rebbeck TR, Dang CV, Zhang L. Integrated Analysis of Genetic Ancestry and Genomic Alterations across Cancers. Cancer Cell. 2018;34(4):549–60 e9. doi: 10.1016/j.ccell.2018.08.019. PubMed PMID: 30300578; PMCID: PMC6348897.

10. Carrot-Zhang J, Chambwe N, Damrauer JS, Knijnenburg TA, Robertson AG, Yau C, Zhou W, Berger AC, Huang KL, Newberg JY, Mashl RJ, Romanel A, Sayaman RW, Demichelis F, Felau I, Frampton GM, Han S, Hoadley KA, Kemal A, Laird PW, Lazar AJ, L. X, Oak N, Shen H, Wong CK, Zenklusen JC, Ziv E, Cancer Genome Atlas Analysis N, Cherniack AD, Beroukhim R. Comprehensive Analysis of Genetic Ancestry and Its Molecular Correlates in Cancer. Cancer Cell. 2020;37(5):639–54 e6. doi: 10.1016/j.ccell.2020.04.012. PubMed PMID: 32396860; PMCID: PMC7328015.

11. DeSantis CE, Miller KD, Goding Sauer A, Jemal A, Siegel RL. Cancer statistics for African Americans, 2019. CA Cancer J Clin. 2019;69(3):211–33. Epub 20190214. doi: 10.3322/caac.21555. PubMed PMID: 30762872.

12. Huo D, Hu H, Rhie SK, Gamazon ER, Cherniack AD, Liu J, Yoshimatsu TF, Pitt JJ, Hoadley KA, Troester M, Ru Y, Lichtenberg T, Sturtz LA, Shelley CS, Benz CC, Mills GB, Laird PW, Shriver CD, Perou CM, Olopade OI. Comparison of Breast Cancer Molecular Features and Survival by African and European Ancestry in The Cancer Genome Atlas. JAMA Oncol. 2017;3(12):1654–62. doi: 10.1001/jamaoncol.2017.0595. PubMed PMID: 28472234; PMCID: PMC5671371.

13. John EM, Phipps AI, Davis A, Koo J. Migration history, acculturation, and breast cancer risk in Hispanic women. Cancer Epidemiol Biomarkers Prev. 2005;14(12):2905–13. doi: 10.1158/1055-9965.EPI-05-0483. PubMed PMID: 16365008.

14. Fejerman L, Ahmadiyeh N, Hu D, Huntsman S, Beckman KB, Caswell JL, Tsung K, John EM, Torres-Mejia G, Carvajal-Carmona L, Echeverry MM, Tuazon AM, Ramirez C, Consortium C, Gignoux CR, Eng C, Gonzalez-Burchard E, Henderson B, Le Marchand L, Kooperberg C, Hou L, Agalliu I, Kraft P, Lindstrom S, Perez-Stable EJ, Haiman CA, Ziv E. Genome-wide association study of breast cancer in Latinas identifies novel protective variants on 6q25. Nat Commun. 2014;5:5260. Epub 20141020. doi: 10.1038/ncomms6260. PubMed PMID: 25327703; PMCID: PMC4204111.

15. Hendrick RE, Monticciolo DL, Biggs KW, Malak SF. Age distributions of breast cancer diagnosis and mortality by race and ethnicity in US women. Cancer. 2021;127(23):4384–92. Epub 20210824. doi: 10.1002/cncr.33846. PubMed PMID: 34427920.

16. Primm KM, Zhao H, Hernandez DC, Chang S. A Contemporary Analysis of Racial and Ethnic Disparities in Diagnosis of Early-Stage Breast Cancer and Stage-Specific Survival by Molecular Subtype. Cancer Epidemiol Biomarkers Prev. 2022;31(6):1185–94. doi: 10.1158/1055-9965.EPI-22-0020. PubMed PMID: 35314859.

17. Fejerman L, Hu D, Huntsman S, John EM, Stern MC, Haiman CA, Perez-Stable EJ, Ziv E. Genetic ancestry and risk of mortality among U.S. Latinas with breast cancer. Cancer Res. 2013;73(24):7243–53. Epub 20131031. doi: 10.1158/0008-5472.CAN-13-2014. PubMed PMID: 24177181; PMCID: PMC3881587.

18. Marker KM, Zavala VA, Vidaurre T, Lott PC, Vasquez JN, Casavilca-Zambrano S, Calderon M, Abugattas JE, Gomez HL, Fuentes HA, Picoaga RL, Cotrina JM, Neciosup SP, Castaneda CA, Morante Z, Valencia F, Torres J, Echeverry M, Bohorquez ME, Polanco-Echeverry G, Estrada-Florez AP, Serrano-Gomez SJ, Carmona-Valencia JA, Alvarado-Cabrero I, Sanabria-Salas MC, Velez A, Donado J, Song S, Cherry D, Tamayo LI, Huntsman S, Hu D, Ruiz-Cordero R, Balassanian R, Ziv E, Zabaleta J, Carvajal-Carmona L, Fejerman L, Consortium C. Human Epidermal Growth Factor Receptor 2-Positive Breast Cancer Is Associated with Indigenous American Ancestry in Latin American Women. Cancer Res. 2020;80(9):1893–901. Epub 20200403. doi: 10.1158/0008-5472.CAN-19-3659. PubMed PMID: 32245796; PMCID: PMC7202960.

19. Romero-Cordoba SL, Salido-Guadarrama I, Rebollar-Vega R, Bautista-Pina V, Dominguez-Reyes C, Tenorio-Torres A, Villegas-Carlos F, Fernandez-Lopez JC, Uribe-Figueroa L, Alfaro-Ruiz L, Hidalgo-Miranda A. Comprehensive omic characterization of breast cancer in Mexican-Hispanic women. Nat Commun. 2021;12(1):2245. Epub 20210414. doi: 10.1038/s41467-021-22478-5. PubMed PMID: 33854067; PMCID: PMC8046804.

20. Chen Y, Lun AT, Smyth GK. From reads to genes to pathways: differential expression analysis of RNA-Seq experiments using Rsubread and the edgeR quasi-likelihood pipeline. F1000Res. 2016;5:1438. Epub 20160620. doi: 10.12688/f1000research.8987.2. PubMed PMID: 27508061; PMCID: PMC4934518.

21. Zhao X, Rodland EA, Tibshirani R, Plevritis S. Molecular subtyping for clinically defined breast cancer subgroups. Breast Cancer Res. 2015;17:29. Epub 20150226. doi: 10.1186/s13058-015-0520-4. PubMed PMID: 25849221; PMCID: PMC4365540.

22. Li R, Qu H, Wang S, Wei J, Zhang L, Ma R, Lu J, Zhu J, Zhong WD, Jia Z. GDCRNATools: an R/Bioconductor package for integrative analysis of lncRNA, miRNA and mRNA data in GDC. Bioinformatics. 2018;34(14):2515–7. doi: 10.1093/bioinformatics/bty124. PubMed PMID: 29509844.

23. Richards S, Aziz N, Bale S, Bick D, Das S, Gastier-Foster J, Grody WW, Hegde M, Lyon E, Spector E, Voelkerding K, Rehm HL, Committee Alqa. Standards and guidelines for the interpretation of sequence variants: a joint consensus recommendation of the American College of Medical Genetics and Genomics and the Association for Molecular Pathology. Genet Med. 2015;17(5):405–24. Epub 20150305. doi: 10.1038/gim.2015.30. PubMed PMID: 25741868; PMCID: PMC4544753.

24. Landrum MJ, Lee JM, Benson M, Brown GR, Chao C, Chitipiralla S, Gu B, Hart J, Hoffman D, Jang W, Karapetyan K, Katz K, Liu C, Maddipatla Z, Malheiro A, McDaniel K, Ovetsky M, Riley G, Zhou G, Holmes JB, Kattman BL, Maglott DR. ClinVar: improving access to variant interpretations and supporting evidence. Nucleic Acids Res. 2018;46(D1):D1062–D7. doi: 10.1093/nar/gkx1153. PubMed PMID: 29165669; PMCID: PMC5753237.

25. Spear ML, Hu D, Pino-Yanes M, Huntsman S, Eng C, Levin AM, Ortega VE, White MJ, McGarry ME, Thakur N, Galanter J, Mak ACY, Oh SS, Ampleford E, Peters SP, Davis A, Kumar R, Farber HJ, Meade K, Avila PC, Serebrisky D, Lenoir MA, Brigino-Buenaventura E, Cintron WR, Thyne SM, Rodriguez-Santana JR, Ford JG, Chapela R, Estrada AM, Sandoval K, Seibold MA, Winkler CA, Bleecker ER, Myers DA, Williams LK, Hernandez RD, Torgerson DG, Burchard EG. A genome-wide association and admixture mapping study of bronchodilator drug response in African Americans with asthma. Pharmacogenomics J. 2019;19(3):249–59. Epub 20180912. doi: 10.1038/s41397-018-0042-4. PubMed PMID: 30206298; PMCID: PMC6414286.

26. Alexander DH, Lange K. Enhancements to the ADMIXTURE algorithm for individual ancestry estimation. BMC Bioinformatics. 2011;12:246. Epub 20110618. doi: 10.1186/1471-2105-12-246. PubMed PMID: 21682921; PMCID: PMC3146885.

27. Chang CC, Chow CC, Tellier LC, Vattikuti S, Purcell SM, Lee JJ. Second-generation PLINK: rising to the challenge of larger and richer datasets. Gigascience. 2015;4:7. Epub 20150225. doi: 10.1186/s13742-015-0047-8. PubMed PMID: 25722852; PMCID: PMC4342193.

28. Van der Auwera GA, O’Connor BD. enomics in the Cloud: Using Docker, GATK, and WDL in Terra (1st Edition). O’Reilly Media; 2020.

29. Tate JG, Bamford S, Jubb HC, Sondka Z, Beare DM, Bindal N, Boutselakis H, Cole CG, Creatore C, Dawson E, Fish P, Harsha B, Hathaway C, Jupe SC, Kok CY, Noble K, Ponting L, Ramshaw CC, Rye CE, Speedy HE, Stefancsik R, Thompson SL, Wang S, Ward S, Campbell PJ, Forbes SA. COSMIC: the Catalogue Of Somatic Mutations In Cancer. Nucleic Acids Res. 2019;47(D1):D941–D7. doi: 10.1093/nar/gky1015. PubMed PMID: 30371878; PMCID: PMC6323903.

30. Lawrence MS, Stojanov P, Polak P, Kryukov GV, Cibulskis K, Sivachenko A, Carter SL, Stewart C, Mermel CH, Roberts SA, Kiezun A, Hammerman PS, McKenna A, Drier Y, Zou L, Ramos AH, Pugh TJ, Stransky N, Helman E, Kim J, Sougnez C, Ambrogio L, Nickerson E, Shefler E, Cortes ML, Auclair D, Saksena G, Voet D, Noble M, DiCara D, Lin P, Lichtenstein L, Heiman DI, Fennell T, Imielinski M, Hernandez B, Hodis E, Baca S, Dulak AM, Lohr J, Landau DA, Wu CJ, Melendez-Zajgla J, Hidalgo-Miranda A, Koren A, McCarroll SA, Mora J, Crompton B, Onofrio R, Parkin M, Winckler W, Ardlie K, Gabriel SB, Roberts CWM, Biegel JA, Stegmaier K, Bass AJ, Garraway LA, Meyerson M, Golub TR, Gordenin DA, Sunyaev S, Lander ES, Getz G. Mutational heterogeneity in cancer and the search for new cancer-associated genes. Nature. 2013;499(7457):214–8. Epub 20130616. doi: 10.1038/nature12213. PubMed PMID: 23770567; PMCID: PMC3919509.

31. Shen R, Seshan VE. FACETS: allele-specific copy number and clonal heterogeneity analysis tool for high-throughput DNA sequencing. Nucleic Acids Res. 2016;44(16):e131. Epub 20160607. doi: 10.1093/nar/gkw520. PubMed PMID: 27270079; PMCID: PMC5027494.

32. Mermel CH, Schumacher SE, Hill B, Meyerson ML, Beroukhim R, Getz G. GISTIC2.0 facilitates sensitive and confident localization of the targets of focal somatic copy-number alteration in human cancers. Genome Biol. 2011;12(4):R41. Epub 20110428. doi: 10.1186/gb-2011-12-4-r41. PubMed PMID: 21527027; PMCID: PMC3218867.

33. Ciriello G, Gatza ML, Beck AH, Wilkerson MD, Rhie SK, Pastore A, Zhang H, McLellan M, Yau C, Kandoth C, Bowlby R, Shen H, Hayat S, Fieldhouse R, Lester SC, Tse GM, Factor RE, Collins LC, Allison KH, Chen YY, Jensen K, Johnson NB, Oesterreich S, Mills GB, Cherniack AD, Robertson G, Benz C, Sander C, Laird PW, Hoadley KA, King TA, Network TR, Perou CM. Comprehensive Molecular Portraits of Invasive Lobular Breast Cancer. Cell. 2015;163(2):506–19. doi: 10.1016/j.cell.2015.09.033. PubMed PMID: 26451490; PMCID: PMC4603750.

34. Blokzijl F, Janssen R, van Boxtel R, Cuppen E. MutationalPatterns: comprehensive genome-wide analysis of mutational processes. Genome Med. 2018;10(1):33. Epub 20180425. doi: 10.1186/s13073-018-0539-0. PubMed PMID: 29695279; PMCID: PMC5922316.

35. Weitzel JN, Neuhausen SL, Adamson A, Tao S, Ricker C, Maoz A, Rosenblatt M, Nehoray B, Sand S, Steele L, Unzeitig G, Feldman N, Blanco AM, Hu D, Huntsman S, Castillo D, Haiman C, Slavin T, Ziv E. Pathogenic and likely pathogenic variants in PALB2, CHEK2, and other known breast cancer susceptibility genes among 1054 BRCA-negative Hispanics with breast cancer. Cancer. 2019;125(16):2829–36. Epub 20190617. doi: 10.1002/cncr.32083. PubMed PMID: 31206626; PMCID: PMC7376605.

36. Lorenzato A, Olivero M, Patane S, Rosso E, Oliaro A, Comoglio PM, Di Renzo MF. Novel somatic mutations of the MET oncogene in human carcinoma metastases activating cell motility and invasion. Cancer Res. 2002;62(23):7025–30. PubMed PMID: 12460923.

37. Alexandrov LB, Kim J, Haradhvala NJ, Huang MN, Tian Ng AW, Wu Y, Boot A, Covington KR, Gordenin DA, Bergstrom EN, Islam SMA, Lopez-Bigas N, Klimczak LJ, McPherson JR, Morganella S, Sabarinathan R, Wheeler DA, Mustonen V, Group Pmsw, Getz G, Rozen SG, Stratton MR, Consortium P. The repertoire of mutational signatures in human cancer. Nature. 2020;578(7793):94–101. Epub 20200205. doi: 10.1038/s41586-020-1943-3. PubMed PMID: 32025018; PMCID: PMC7054213.

38. Kanu N, Cerone MA, Goh G, Zalmas LP, Bartkova J, Dietzen M, McGranahan N, Rogers R, Law EK, Gromova I, Kschischo M, Walton MI, Rossanese OW, Bartek J, Harris RS, Venkatesan S, Swanton C. DNA replication stress mediates APOBEC3 family mutagenesis in breast cancer. Genome Biol. 2016;17(1):185. Epub 20160915. doi: 10.1186/s13059-016-1042-9. PubMed PMID: 27634334; PMCID: PMC5025597.

39. Komatsu A, Nagasaki K, Fujimori M, Amano J, Miki Y. Identification of novel deletion polymorphisms in breast cancer. Int J Oncol. 2008;33(2):261–70. PubMed PMID: 18636146.

40. Caval V, Suspene R, Shapira M, Vartanian JP, Wain-Hobson S. A prevalent cancer susceptibility APOBEC3A hybrid allele bearing APOBEC3B 3’UTR enhances chromosomal DNA damage. Nat Commun. 2014;5:5129. Epub 20141009. doi: 10.1038/ncomms6129. PubMed PMID: 25298230.

41. Xuan D, Li G, Cai Q, Deming-Halverson S, Shrubsole MJ, Shu XO, Kelley MC, Zheng W, Long J. APOBEC3 deletion polymorphism is associated with breast cancer risk among women of European ancestry. Carcinogenesis. 2013;34(10):2240–3. Epub 20130528. doi: 10.1093/carcin/bgt185. PubMed PMID: 23715497; PMCID: PMC3786378.

42. Nik-Zainal S, Davies H, Staaf J, Ramakrishna M, Glodzik D, Zou X, Martincorena I, Alexandrov LB, Martin S, Wedge DC, Van Loo P, Ju YS, Smid M, Brinkman AB, Morganella S, Aure MR, Lingjaerde OC, Langerod A, Ringner M, Ahn SM, Boyault S, Brock JE, Broeks A, Butler A, Desmedt C, Dirix L, Dronov S, Fatima A, Foekens JA, Gerstung M, Hooijer GK, Jang SJ, Jones DR, Kim HY, King TA, Krishnamurthy S, Lee HJ, Lee JY, Li Y, McLaren S, Menzies A, Mustonen V, O’Meara S, Pauporte I, Pivot X, Purdie CA, Raine K, Ramakrishnan K, Rodriguez-Gonzalez FG, Romieu G, Sieuwerts AM, Simpson PT, Shepherd R, Stebbings L, Stefansson OA, Teague J, Tommasi S, Treilleux I, Van den Eynden GG, Vermeulen P, Vincent-Salomon A, Yates L, Caldas C, van’t Veer L, Tutt A, Knappskog S, Tan BK, Jonkers J, Borg A, Ueno NT, Sotiriou C, Viari A, Futreal PA, Campbell PJ, Span PN, Van Laere S, Lakhani SR, Eyfjord JE, Thompson AM, Birney E, Stunnenberg HG, van de Vijver MJ, Martens JW, Borresen-Dale AL, Richardson AL, Kong G, Thomas G, Stratton MR. Landscape of somatic mutations in 560 breast cancer whole-genome sequences. Nature. 2016;534(7605):47–54. Epub 20160502. doi: 10.1038/nature17676. PubMed PMID: 27135926; PMCID: PMC4910866.

43. Song J, Yang W, Shih Ie M, Zhang Z, Bai J. Identification of BCOX1, a novel gene overexpressed in breast cancer. Biochim Biophys Acta. 2006;1760(1):62–9. Epub 20051025. doi: 10.1016/j.bbagen.2005.09.017. PubMed PMID: 16289875.

44. Liu T, Zhang XY, He XH, Geng JS, Liu Y, Kong DJ, Shi QY, Liu F, Wei W, Pang D. High levels of BCOX1 expression are associated with poor prognosis in patients with invasive ductal carcinomas of the breast. PLoS One. 2014;9(1):e86952. Epub 20140128. doi: 10.1371/journal.pone.0086952. PubMed PMID: 24489812; PMCID: PMC3904964.

45. Zhong Z, Pannu V, Rosenow M, Stark A, Spetzler D. KIAA0100 Modulates Cancer Cell Aggression Behavior of MDA-MB-231 through Microtubule and Heat Shock Proteins. Cancers (Basel). 2018;10(6). Epub 20180604. doi: 10.3390/cancers10060180. PubMed PMID: 29867023; PMCID: PMC6025110.

46. Thompson AM, Moulder-Thompson SL. Neoadjuvant treatment of breast cancer. Ann Oncol. 2012;23 Suppl 10:x231–6. doi: 10.1093/annonc/mds324. PubMed PMID: 22987968; PMCID: PMC6278992.

47. Zavala VA, Bracci PM, Carethers JM, Carvajal-Carmona L, Coggins NB, Cruz-Correa MR, Davis M, de Smith AJ, Dutil J, Figueiredo JC, Fox R, Graves KD, Gomez SL, Llera A, Neuhausen SL, Newman L, Nguyen T, Palmer JR, Palmer NR, Perez-Stable EJ, Piawah S, Rodriquez EJ, Sanabria-Salas MC, Schmit SL, Serrano-Gomez SJ, Stern MC, Weitzel J, Yang JJ, Zabaleta J, Ziv E, Fejerman L. Cancer health disparities in racial/ethnic minorities in the United States. Br J Cancer. 2021;124(2):315–32. Epub 20200909. doi: 10.1038/s41416-020-01038-6. PubMed PMID: 32901135; PMCID: PMC7852513.

